# Integrin binding dynamics modulate ligand-specific mechanosensing in mammary gland fibroblasts

**DOI:** 10.1101/570721

**Authors:** Martina Lerche, Alberto Elosegui-Artola, Jenny Z. Kechagia, Camilo Guzmán, Maria Georgiadou, Donald Gullberg, Pere Roca-Cusachs, Emilia Peuhu, Johanna Ivaska

## Abstract

The link between the modulation of integrin activity and cellular mechanosensing of tissue rigidity, especially on different extracellular matrix ligands, remains poorly understood. Here, we find that primary mouse mammary gland stromal fibroblasts (MSFs) are able to spread efficiently on soft collagen-coated substrates, resembling the soft mammary gland tissue. In addition, MSFs generate high forces and display nuclear YAP at a low matrix stiffness, supported by mature focal adhesions, prominent actin stress fibers, and myosin phosphorylation.

We describe that loss of the cytosolic integrin inhibitor, SHARPIN, impedes MSF spreading specifically on soft type I collagen but not on fibronectin. Through quantitative experiments and computational modelling, we find that SHARPIN-deficient MSFs display faster force-induced unbinding of adhesions from collagen-coated beads. Faster unbinding, in turn, impairs force transmission in these cells, particularly, at the stiffness optimum observed for wild-type cells, and increases actin retrograde flow. Mechanistically, we link the impaired mechanotransduction of SHARPIN-deficient cells on collagen to reduced levels of the collagen-binding integrin α11β1. Our results unveil a collagen-specific mechanosensing mechanism and suggest a key function for integrin activity regulation and integrin α11β1 in MSF mechanotransduction.

## Introduction

Fibroblasts exert high forces that are implicated in the morphogenetic rearrangement of extracellular matrices (ECMs) (Harris et al., 1981). In the developing mammary gland, stromal cell-mediated organization of the ECM regulates mammary ductal morphogenesis (Brownfield et al., 2013; Ingman et al., 2006). Despite this important function, investigations into mammary stromal components are secondary to that of the mammary epithelium. In addition to ECM organization, mammary gland stromal fibroblasts (MSFs) also play a central role in the pro-invasive stiffening of breast tumor stroma (Navab et al., 2016), and therefore understanding the mechanical aspects of these cells is of clinical interest. While the role of integrins as cell mechanosensors and transducers is well established, the link between the regulation of integrin activity and the mechanosensing response on different ECM ligands remains poorly understood. SHARPIN is a cytosolic adaptor protein that, among other functions, binds to the intracellular integrin alpha tails and inhibits integrin activity in different cell types *in vitro* and *in vivo* (Kasirer-Friede et al., 2019; Peuhu, Kaukonen et al., 2017; Peuhu, Salomaa et al., 2017; Pouwels et al., 2013; Rantala et al., 2011). We have previously demonstrated that stromal SHARPIN deficiency interferes with normal mouse mammary gland development and collagen fiber assembly *in vivo* (Peuhu, Kaukonen et al., 2017). However, how SHARPIN mediates integrin-dependent mechanotransduction remains unresolved.

Collagen is abundant in the mammary gland stroma and plays a key role in regulating the physical and biochemical properties of the mammary gland. Alignment of stromal collagen bundles is critical for normal mammary gland development providing migration cues to the outgrowing duct during puberty (Brownfield et al., 2013; Ingman et al., 2006). There are four collagen-binding integrin heterodimers in mammals: The more ubiquitously expressed α1β1, α2β1, and α11β1 and the cartilage specific α10β1 (Zeltz and Gullberg, 2016). Of these, the fibrillar collagen-binding integrins α2β1 and α11β1 have been strongly linked to collagen remodeling and turnover (Abair et al., 2008; Ivaska et al., 1999; Popova et al., 2007; Riikonen et al., 1995; Tiger et al., 2001) and α11β1 to the induction of cancer stromal stiffness (Navab et al., 2016; Zeltz et al., 2019). Furthermore, “trail blazer” breast cancer cells with high invasive capacity are characterized by high integrin α11β1 expression (Westcott et al., 2015). Nevertheless, integrin α11β1 functions are rather poorly understood, and the role of this receptor in regulating cell-collagen interactions in the mammary gland has not been previously studied.

In order to sense the properties of the surrounding ECM, cells use dynamic molecular bonds, often referred to as molecular clutches, to exert forces within the cell boundary (Elosegui-Artola et al., 2018). A molecular clutch can be defined as a dynamic link between the ECM, integrin adhesion receptors, intracellular adaptor proteins, and the actomyosin cytoskeleton (Elosegui-Artola et al., 2014; Elosegui-Artola et al., 2016). By quantification of the molecular clutch binding dynamics, and using mathematical modelling, one can predict the average force transmission of cells to the ECM as a function of substrate stiffness (Elosegui-Artola et al., 2014; Elosegui-Artola et al., 2016).

Here, we have combined mathematical modelling with cell biology to investigate the biomechanical properties of primary mouse MSFs and to understand how the integrin inhibitor SHARPIN affects integrin-dependent force generation and mechanotransduction. We find that, somewhat counterintuitively, in spite of having higher integrin β1 activity SHARPIN-deficient MSFs were defective in spreading on soft hydrogels with a stiffness similar to the mammary gland tissue *in vivo.*They also had faster force induced cell-ECM unbinding rates. Interestingly, both of these defects were specific to collagen and not observed on fibronectin. The molecular clutch model predicted that increased clutch unbinding rates result in the loss of stiffness-dependent traction maximum and increased actin flow rates at low rigidities. Importantly, these predictions were recapitulated experimentally in SHARPIN-deficient primary MSFs on collagen I. SHARPIN-deficient MSFs had significantly downregulated collagen-binding integrin α11β1 levels, explaining mechanistically their unexpected inability to couple to collagen. These data highlight an important divergence in the regulation of collagen I-and fibronectin-binding integrin heterodimers in the mammary gland stroma with implications for the mechanical response of fibroblasts. Moreover, these insights are likely to improve our understanding of fibrotic diseases including cancer where fibroblasts exhibit deregulated integrin activity (Erdogan et al., 2017; Glentis et al., 2017).

## Results

### Increased integrin activity correlates with reduced spreading of mouse mammary gland fibroblasts on soft collagen I-but not on soft fibronectin-coated matrices

Based on previous observations that primary MSFs from SHARPIN-deficient mice (chronic proliferative dermatitis null mutation, *cpdm; Sharpin^cpdm/cpdm^; Sharpin^cpdm^)(HogenEsch* et al., 1993; Seymour et al., 2007) have increased integrin β1 activity but, counterintuitively, impaired capacity to contract collagen gels (Peuhu, Kaukonen et al., 2017) we sought to investigate the link between integrin activity and force transduction. We first confirmed by flow cytometry the cell-surface levels of total and active integrin β1 with conformation-specific antibodies. As expected, *Sharpin^cpdm^* MSFs expressed lower total integrin β1 cell-surface levels but equal levels of active integrin β1 compared to SHARPIN-expressing *(Sharpin^+/+^or Sharpin^cpdm/+^;* from here on referred to as wild-type) cells, indicating that in *Sharpin^cpdm^* MSFs a higher proportion of integrin β1 is in the active conformation on the cell surface (Fig. 1A, S1A), in line with our previous studies with MSFs (Peuhu, Kaukonen et al., 2017) and other cell types (Peuhu, Salomaa et al., 2017; Rantala et al., 2011). Next, we studied the ability of wild-type and *Sharpin^cpdm^* MSFs to spread in response to ECM stiffness and ligand type. MSFs were seeded at equal density on soft (2 kPa) fibronectin or collagen I pre-coated polyacrylamide gels, approximating the stiffness of the mammary tissue *in vivo* (Lopez et al., 2011; Peuhu, Kaukonen et al., 2017; Plodinec et al., 2012). As expected based on the higher integrin β1 activity and faster focal adhesion (FA) turnover compared to wild-type MSFs (Peuhu, Kaukonen et al., 2017; Rantala et al., 2011), *Sharpin^cpdm^* MSFs spread more compared to wild-type MSFs when seeded on fibronectin-coated hydrogels (Fig. 1B, C). In contrast, on 2 kPa collagen I-coated hydrogels *Sharpin^cpdm^* MSFs were less spread than wild-type MSFs (Fig. 1B, C). When cell spreading area was measured on a stiffness range from 0.8-10 kPa (Fig. 1C), on collagen I-coated hydrogels, wild-type MSFs displayed a spreading optimum on 2 kPa, while *Sharpin^cpdm^* MSFs were significantly smaller and only fully spread at 10 kPa (Fig. 1C). These data present an unexpected conundrum; at lower stiffness (corresponding to the higher end of the rigidity spectrum reported for mammary gland tissue *in vivo*, 0.1-2 kPa (Peuhu, Kaukonen et al., 2017; Plodinec et al., 2012), loss of SHARPIN (coinciding with increased integrin β1 activity) correlates with defective MSF spreading on collagen I, while on fibronectin the opposite is observed.

**Figure 1.**
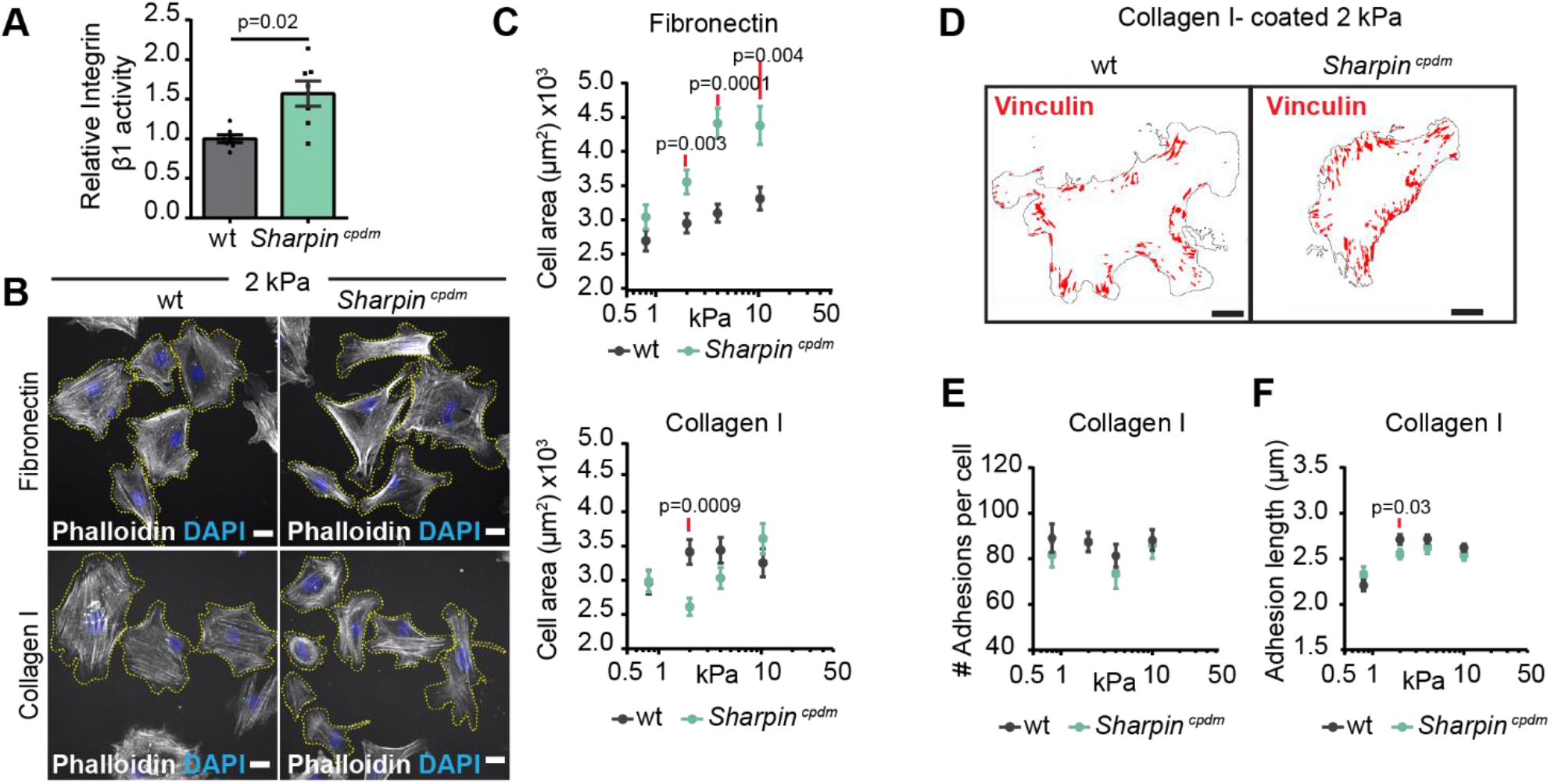
Increased integrin activity correlates with reduced spreading of MSFs on soft collagen I. (**A**) Quantification of relative integrin β1 activity (active (clone, 9EG7) / total (clone, HMβ1-1)) cellsurface levels (n=7 independent experiments) in *Sharpin^cpdm^* compared to wild-type MSFs by flow cytometry. (**B**) Representative images of wild-type and *Sharpin^cpdm^* MSFs plated for 3-4 h on 2 kPa fibronectin (upper panel) or collagen I (lower panel)-coated polyacrylamide (PAA) hydrogels and labelled for F-actin (white) and nuclei (blue). Cell edges are outlined with yellow dashed lines. (**C**) Quantification of cell spreading in wild-type compared to *Sharpin^cpdm^* MSFs on 0.8, 2, 4 and 13 kPa PAA hydrogels coated with fibronectin (upper panel) or collagen I (lower panel) based on immunofluorescence. Data are pooled from 3 independent experiments, n_wt_= 90, 95, 103, 88 and n_Sharpincpdm_= 72, 80, 93, 88 cells (fibronectin, from left to right) and n_wt_= 64, 105, 91, 109 and n_Sharpincpdm_= 59, 118, 97, 77 cells (collagen I, from left to right). (**D**) Representative output images of FA analysis for individual cells plated on 2 kPa collagen I-coated PAA hydrogels (FA, red; cell borders, black). (**E, F**) Quantification of the number of FA per cell (E) and the length of FA (F) in wild-type compared to *Sharpin^cpdm^* MSFs plated on collagen I-coated 0.8, 2, 4 and 13 kPa PAA hydrogels. Data are pooled from three independent experiments, n_wt_= 64, 94, 73, 83, n_Sharpincpdm_= 59, 97, 73, 56 cells (# Adhesion per cell, from left to right) and n_wt_= 41, 94, 73, 109 and n_Sharpincpdm_= 42, 98, 74, 73 cells (Adhesion length, from left to right). Mean ± SEM in all graphs. Mann–Whitney *U*-test; red lines above p-values indicate the data points compared. Scale bars: 20 μm.

Given that *Sharpin^cpdm^* MSFs have faster FA dynamics (increased assembly and disassembly rates) on collagen-coated rigid glass substrate (Peuhu, Kaukonen et al., 2017), we next evaluated the effect of SHARPIN deficiency on FA maturation (number and average length) on a range of hydrogel rigidities. Vinculin, a mechanosensitive adaptor molecule recruited to mature FA (Chen et al., 2006; del Rio et al., 2009), was immunolabeled in wild-type and *Sharpin^cpdm^* MSFs plated on 0.8-10 kPa collagen I-coated hydrogels (Fig. 1D, Fig. S1B). Wild-type and *Sharpin^cpdm^* MSFs (Fig. 1D) demonstrated similar FA number and length on collagen I-coated hydrogels (Fig. 1E, F). Interestingly, both wild-type and *Sharpin^cpdm^* primary MSFs formed mature, vinculin-containing adhesions on soft matrices reaching the maturation maxima already on 2-4 kPa (Fig 1F). On fibronectin, *Sharpin^cpdm^* MSFs demonstrated either longer or more FA on 2-4 kPa hydrogels (Fig. S1C-E), consistent with increased cell spreading (Fig. 1B, C). Furthermore, MSFs exhibited nuclear localization of the mechanosensitive transcription factor Yes-associated protein (YAP) at 2 kPa stiffness (Fig. S1F). This localization was reduced in *Sharpin^cpdm^* MSFs (Fig. S1F, G). The nuclear localization of YAP and the ability of MSFs to generate elongated, stress-fiber linked adhesions on a low stiffness is counter to the stiffness induced adhesion reinforcement/maturation and nucler YAP translocation detected in other cell types (Elosegui-Artola et al., 2016), possibly mirroring the adaption of these primary cells to their soft growth environment in the mammary gland (Lopez et al., 2011; Peuhu, Kaukonen et al., 2017; Plodinec et al., 2012). In conclusion, *Sharpin^cpdm^* MSFs demonstrate reduced capacity to spread on soft collagen I-coated matrices, while the opposite occurs on fibronectin-coated substratum.

### Increased integrin activity correlates with faster integrin-collagen binding dynamics in MSFs

We then assessed how the altered integrin activity in *Sharpin^cpdm^* MSFs affects the binding and unbinding properties of the cells to matrix ligands (Fig. 2A). For this we employed ECM-coated bead recruitment and -detachment experiments. Previous studies indicate that adhesions formed by cells on beads of comparable size [3 μm vs. 2.8-4.5 μm; (Guilluy et al., 2011; Jones et al., 2015)], have similar attachment times [30 min vs. 20-35 min; (González-Tarragó et al., 2017; Jones et al., 2015)] and recruit the same core-adhesome proteins as the adhesions that form on the basal side of adherent cells (Guilluy et al., 2011; Jones et al., 2015). *Sharpin^cpdm^* MSFs displayed slightly increased recruitment of integrin β1 to collagen I-coated silica beads when compared to wild-type cells, while no significant differences in binding to fibronectin-coated beads were observed (Fig. 2B). We then employed a magnetic tweezers setup (Elosegui-Artola et al., 2014), a method that allows quantitative measurement of the strength of receptor-ligand bonds, to apply force to collagen I or fibronectin-coated beads attached to cells, and evaluated the time required to detach beads from cells (Fig. 2A). Detachment times of collagen-coated beads were significantly lower for *Sharpin^cpdm^* MSFs compared to wild type cells (Fig. 2C). However, no significant differences were observed with fibronectin coated beads (Fig. 2C). The significant decrease in bead detachment time under force suggests that unbinding rates under force are increased. Thus, overall the results are consistent with an effect of SHARPIN deficiency in increasing both binding and unbinding rates of integrins to collagen, thereby maintaining the overall recruitment constant.

**Figure 2.**
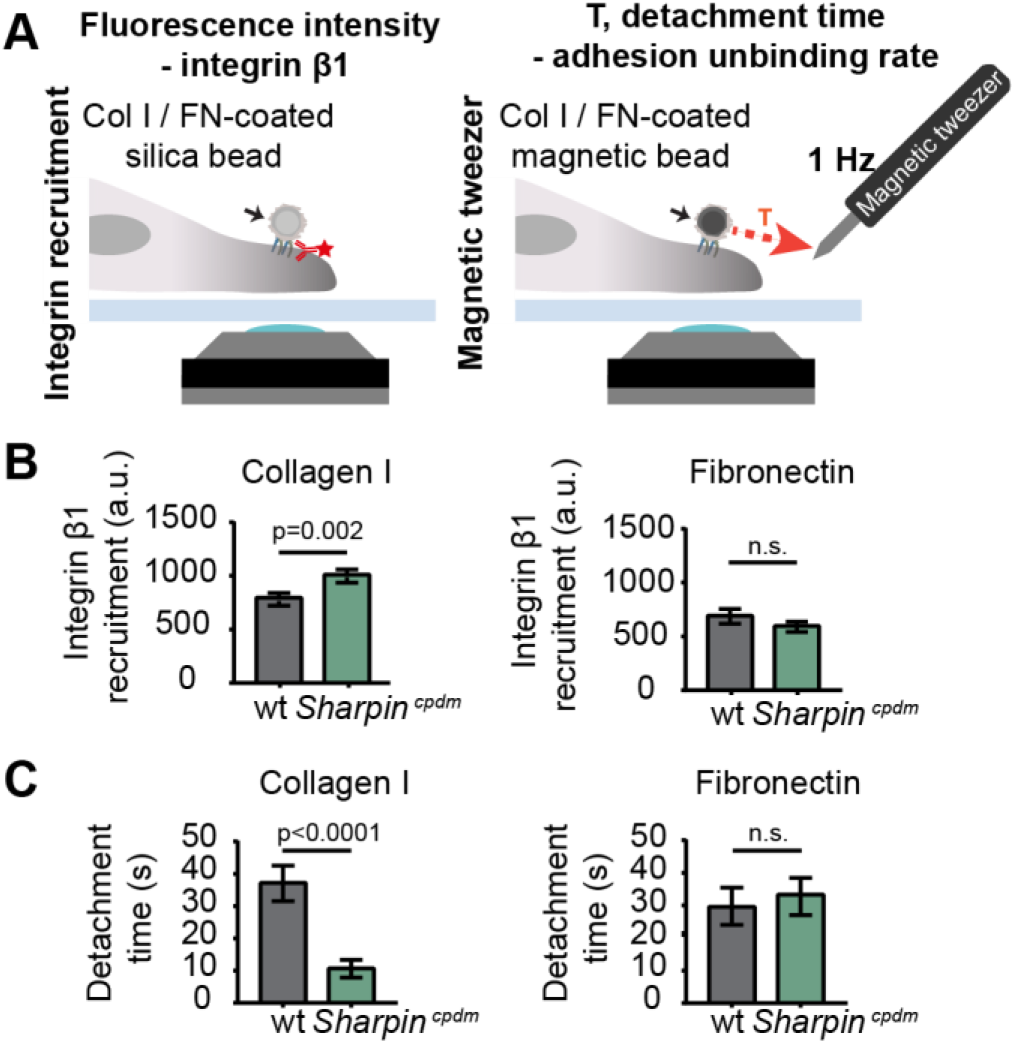
SHARPIN-deficient MSFs show faster integrin-collagen binding dynamics. (**A**) Schematic representation of the set up for integrin recruitment (left panel) and magnetic tweezer (right panel) experiments. (**B**) Quantification of integrin β1 recruitment to collagen I (left panel) or fibronectin (right panel) -coated silica beads; n_wt_=62 and n_Sharpincpdm_=51 cells (collagen I) and n_wt_=33 and n_Sharpincpdm_=24 cells (fibronectin) from 2 independent experiments. (**C**) Quantification of detachment time of wild-type and *Sharpin^cpdm^* MSFs from collagen I (left panel) or fibronectin (right panel) - coated magnetic beads; nwt and n_Sharpincpdm_=34 cells (collagen) and n_wt_=29 and n_Sharpincpdm_=37 cells (fibronectin) from 2 independent experiments. Mean ± SEM in all graphs. Unpaired t-test. Col I, collagen I; FN, fibronectin.

### Molecular clutch model predicts the absence of traction peak in Sharpin^cpdm^ cells at biologically relevant rigidities

The ability of cells to sense and respond to rigidity, and apply force to the matrix, is regulated by the different components of the adhesive and contractile molecular machinery that functions jointly as a cellular molecular clutch (Case and Waterman, 2015; Chan and Odde, 2008; Elosegui-Artola et al., 2014). Clutch dynamics are a function of several parameters: The number and binding dynamics of integrin-ECM bonds, reinforcement of the integrin-actin link through talin unfolding and vinculin recruitment, actomyosin contractility, and substrate compliance. Unlike previously studied cell types (Elosegui-Artola et al., 2016), the MSFs are able to to generate fully mature adhesions at low stiffness. This suggests that MSFs should exhibit the fundamental prediction of the molecular clutch, a biphasic force/rigidity relationship, which is almost always masked by the fact that adhesion growth and reinforcement normally occur only at high rigidities (Chan and Odde, 2008; Elosegui-Artola et al., 2016). This exciting scenario prompted us to measure the remaining necessary parameters required for computational modelling of the molecular clutch in these cells. Actomyosin contractility is an important component of the molecular clutch and is regulated by phosphorylation of myosin light chain 2 (pMLC2). In *talin 1−/−* knockout mouse embryonic fibroblasts (MEFs) (talin 1^−/−^ MEFs), which display a wild-type phenotype due to compensatory upregulation of endogenous talin 2 and have previously been studied with comparable methods (Elosegui-Artola et al., 2016), pMLC2 levels are largely independent of substrate stiffness (Elosegui-Artola et al., 2016). We compared our MSFs to the talin 1^−/−^ MEFs and observed increasing pMLC2 levels in MSFs in response to increasing stiffness (Fig. 3A, Fig. S2A, B), similar to the previously observed response of vascular smooth muscle cells (Polte et al., 2004). Furthermore, higher pMLC was observed in MSFs plated on soft substrate as compared to talin 1 ^−/−^ MEFs (Elosegui-Artola et al., 2016) (Fig. S2A). However, no significant differences in pMLC2 were detected between wild-type and *Sharpin^cpdm^* MSFs, suggesting that myosin activity remains predominantly unaffected in the absence of SHARPIN.

**Figure 3.**
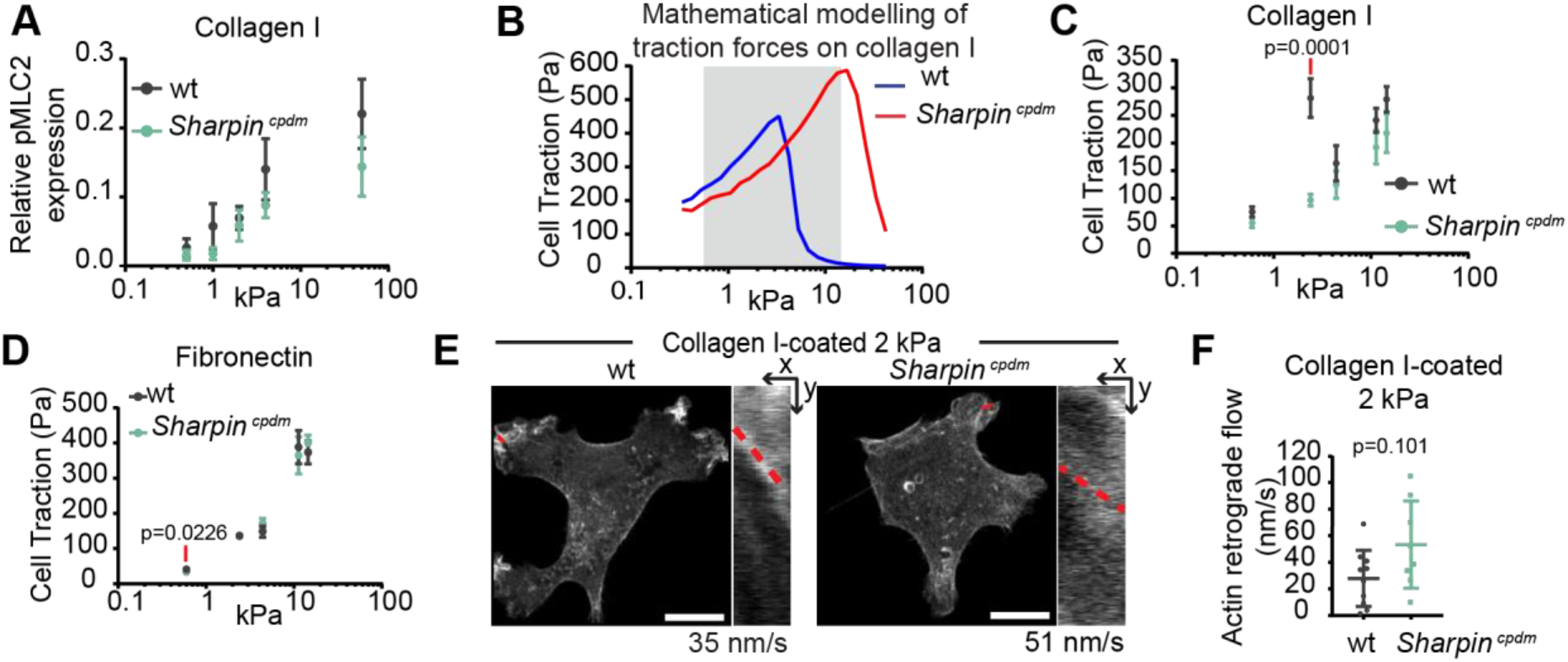
Molecular clutch model predicts the absence of traction peak in SHARPIN-deficient cells at biologically relevant rigidities. (**A**) Quantification of relative pMLC2 expression levels in wild-type compared to *Sharpin^cpdm^* MSFs plated on collagen I-coated PAA hydrogels with the indicated stiffness, n= 3 independent experiments. (**B**) Prediction of the traction forces generated by wildtype and *Sharpin^cpdm^* MSFs on collagen I-coated PAA hydrogels based on the molecular clutch model. The stiffness range covered in (**C**) is highlighted. (**C**) Average forces exerted by wild-type compared to *Sharpin^cpdm^* MSFs on collagen I-coated PAA hydrogels with the indicated stiffness measured by traction force microscopy, n_wt_= 20, 20, 21, 20, 19 and n_Sharpincpdm_= 18, 25, 21, 18, 17 cells (from left to right) from 2 independent experiments. (**D**) Average forces exerted by wild-type compared to *Sharpin^cpdm^* MSFs on fibronectin-coated PAA hydrogels with the indicated stiffness measured by traction force microscopy, n_wt_= 11, 11, 23, 14, 10 and n_Sharpincpdm_= 10, 13, 23, 11, 10 cells (from left to right) from 2 independent experiments. (**E**) Representative images of Lifeact-GFP transfected wildtype and *Sharpin^cpdm^* MSFs plated on 2 kPa collagen I-coated PAA hydrogels. Insets are kymographs showing actin retrograde flow along the red line (time=180s, imaged every second). The slope of the line was used to calculate the actin retrograde flow rate. (**F**) Quantification of actin retrograde flow in wild-type compared to *Sharpin^cpdm^* MSFs, n_wt_=10 and n_Sharpincpdm_=8 cells (1 measurement/cell), from 3 independent experiments. Mean ± SEM in all graphs. Mann–Whitney *U*-test, red lines above p-values indicate the data points compared. Scale bars: 10 μm.

Taking into consideration these experimentally determined MSF features, we employed the computational model of the molecular clutch (Elosegui-Artola et al., 2014; Elosegui-Artola et al., 2016). It predicts that in the absence of changes in integrin recruitment or adhesion growth as a function of stiffness, the cells should display a biphasic force/rigidity response, even if myosin contractility increases with rigidity (Fig. 3B). The other prediction arising from our modelling is that increased integrin binding and unbinding rates, as observed in *Sharpin^cpdm^* MSFs, should displace the traction force peak to higher rigidities. To test the model predictions, we measured cell-matrix force transmission using traction force microscopy. As predicted, wild-type cells on collagen I displayed a biphasic response of force as a function of stiffness, on the lower stiffness range, with a force maximum at 2 kPa (Fig. 3C). Thus, MSFs represent the first mammalian primary cell type that shows this predicted biphasic force response without genetic perturbation. In *Sharpin^cpdm^* cells, the concomitant increase in binding and unbinding rates should shift the force curve, displacing the force peak to higher rigidities. Accordingly, the maximum force peak at 2 kPa was absent (Fig. 3C), closely replicating the result of the molecular clutch modelling (Fig. 3B). In this case, no force peak was observed, potentially because it was displaced to a stiffness above the range where traction forces were measured experimentally. Of note, wild-type cells exhibit a stiffness-dependent additional increase in force transmission above 4 kPa that is not predicted by the model indicating that the modelling corresponds to the experimental data only in the lower stiffness range. The nature of this regime remains unknown and warrants further investigation.

Finally, the molecular clutch model also predicts that changes in force should inversely correlate with actin flow. As predicted, actin flow of actively spreading (plated for 45-105 minutes) *Sharpin^cpdm^* cells (average 51 nm/s) on 2 kPa collagen I-coated hydrogels was also increased with respect to wildtype cells (average 35 nm/s) (Fig. 3E, F). Due to the limited transfection efficiency in the primary MSFs and the need to isolate them freshly from mouse mammary gland, this single stiffness was selected based on the highest differences in traction force measurements. Interestingly, in stably adhered MSFs (plated for 4 hours) on 2 kPa collagen I-coated hydrogels, very slow actin retrograde flow was observed compared to talin 1^−/−^ MEFs (Fig. S2C), and measurement of actin flow in MSFs was beyond the detection limit. This is in stark contrast to the rapid actin flow detected in other cell types on soft substrate (Bangasser et al., 2017; Chan and Odde, 2008; Elosegui-Artola et al., 2016), further demonstrating the adaption of MSFs to a soft environment. This may be attributed, in part, to integrin expression. We compared the cell-surface levels of several integrin subunits in talin 1^−/−^MEFs and wild-type MSFs and observed that MSFs express nearly two times more integrin β1 and markedly more integrin α1- and α11-subunits compared to talin 1^−/−^ MEFs (Fig. S2D) that spread poorly on soft collagen (Fig. S2E).

Consistent with the fact that no differences in adhesion behavior under force were observed with fibronectin-coated beads (Fig. 2C), wild-type and *Sharpin^cpdm^* cells exerted the same forces on fibronectin-coated substrates irrespective of stiffness and the cells displayed monotonic increase in force with increasing rigidity (Fig. 3D). In contrast to collagen I substrate, MSFs did not show a force maximum at low rigidities on fibronectin. As in *Sharpin^cpdm^* cells on collagen, this lack of a biphasic force response could result from the limited stiffness range used in these experiments (0.6-14.5 kPa), and suggests that the force maximum for MSF on fibronectin is exerted at a stiffness above 14.5 kPa. Our observations support the view that different integrin heterodimers form integrin-ligand bonds with different strengths depending also on ECM composition, which leads to variations in the force maximum on different ECM ligands.

Together, these data demonstrate that SHARPIN deficiency, and the consequent increase in integrin-collagen unbinding rate, leads to significant effects in mechanotransduction in MSFs, providing a possible explanation to our previous finding that *Sharpin^cpdm^* MSFs are unable to remodel collagen *in vitro* and are defective in supporting generation of mammary gland stromal architecture supportive of normal development and ductal outgrowth (Peuhu, Kaukonen et al., 2017).

### Integrin α11β1 protein levels regulate the spreading of MSFs on soft matrices

Next, we asked how loss of SHARPIN could affect integrin binding dynamics, and what the consequent effects are. One potential explanation is that *Sharpin^cpdm^* and wild-type MSFs exhibit differences in collagen-binding integrin expression, leading to different ECM binding properties. An RNA sequencing dataset of wild-type and *Sharpin^cpdm^* MSFs (Peuhu, Kaukonen et al., 2017) was analyzed for all the matrix-binding integrin subtypes (Fig. S3A). Of the collagen-binding integrin alpha subunits *(Itga1, Itga2, Itga10, Itga11)*, which all form a heterodimer with the integrin β1 subunit, *Itga11* was the predominant α-subunit expressed at mRNA level, *Itga1* was detected at low levels and *Itga10* or *Itga2* were not detected. Importantly, no significant differences in integrin mRNA expression levels were observed between wild-type and *Sharpin^cpdm^* MSFs (Fig. S3A).

Next we analysed the cell-surface levels of the expressed collagen binding integrin α-subunits. *Sharpin^cpdm^* MSFs had significantly reduced levels of integrins α1 and α11 compared to wild-type MSFs (Fig. 4A). Western blotting and immunofluorescence labeling further confirmed that integrin α11 levels were lower in *Sharpin^cpdm^* MSFs (Fig. 4B, C, D; mouse integrin α1 protein levels were not analyzed with western blotting or immunofluorescence labeling due to the lack of suitable antibodies). In addition, siRNA-mediated downregulation of SHARPIN reduced integrin α11 protein levels compared to control silencing (Fig. 4E, F). These data suggest that integrin α11 is acutely regulated by SHARPIN and the low integrin α11 levels are not triggered by an *in vivo* compensatory effect in the *Sharpin^cpdm^* animals. A reduced protein level of integrin α11 was observed also in *Sharpin^cpdm^* MSFs that were seeded on 2 kPa collagen I-coated hydrogels directly after isolation from the mouse mammary gland (Fig. S3B), indicating that the observed difference in integrin α11 is not induced by *in vitro* culture on stiff fibronectin-rich substratum (plastic in the presence of serum fibronectin) and is prominent under conditions when collagen I is the only provided ligand (Fig. S3C).

**Figure 4.**
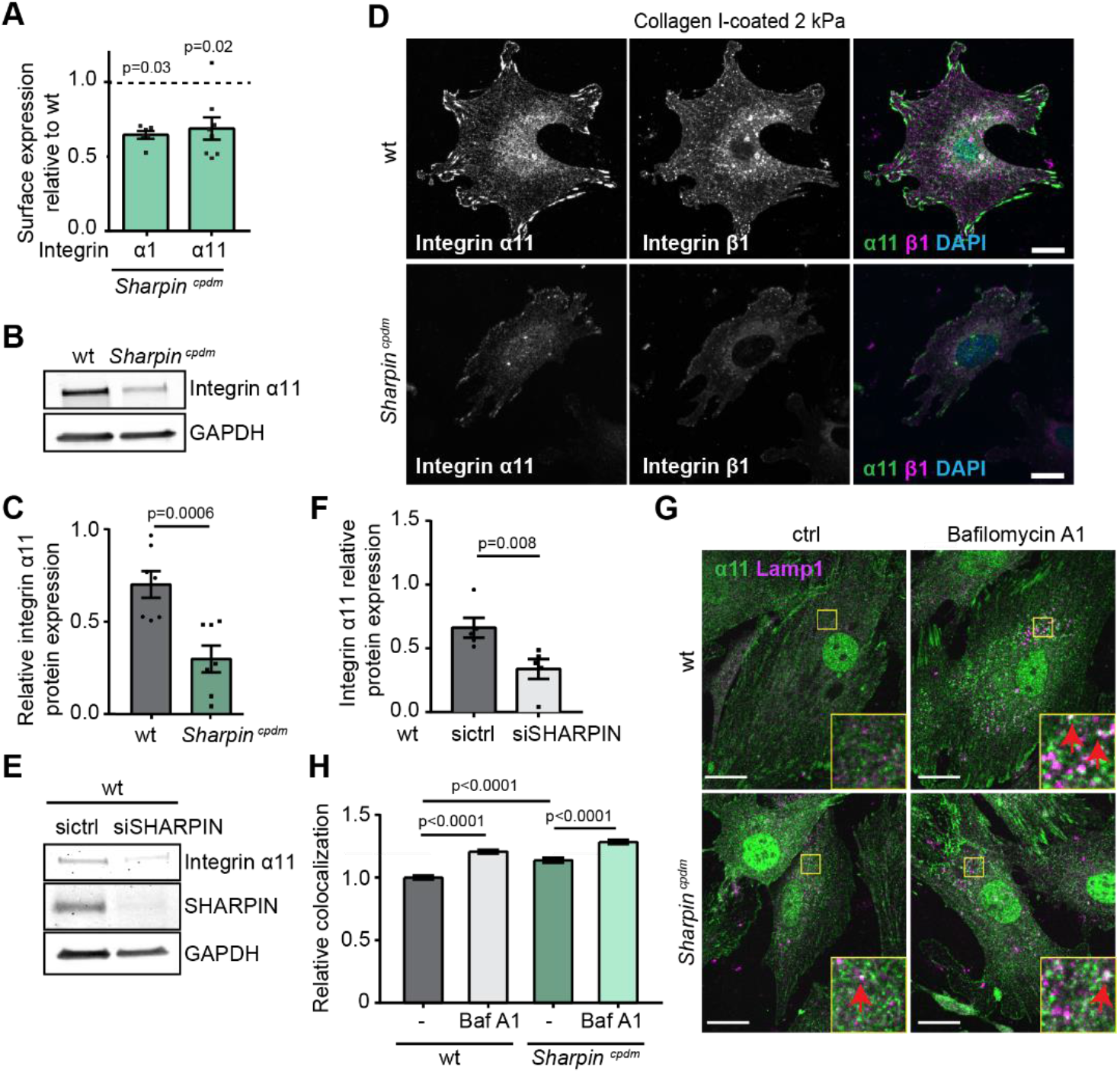
SHARPIN regulates integrin α11β1 protein levels. (**A**) Analysis of cell-surface expression (median fluorescence) of integrin α1 and-α11 in *Sharpin^cpdm^* relative to wild-type MSFs, n_Itga1_ =6 and n_Itga11_=8 from 6 independent flow cytometry experiments. (**B**) Representative Western blot analysis of integrin α11 protein expression in wild-type and *Sharpin^cpdm^* MSFs, and (**C**) quantification of the relative integrin α11 expression levels, nwt and n_Sharpincpdm_=7 from 5 independent experiments. GAPDH was detected for loading control. (**D**) Representative images of immunolabelled integrin α11 (green) and total integrin β1 (magenta) in wild-type and *Sharpin^cpdm^* MSFs plated on 2 kPa collagen I-coated PAA hydrogels. Nuclei (blue) were co-labelled. (**E**) Representative Western blot analysis of integrin α11 and SHARPIN protein expression in wild-type MSFs silenced with control or SHARPIN-targeting siRNA, and (**F**) quantification of the relative integrin α11 expression levels n=5 independent experiments. GAPDH was detected for loading control. (**G**) Representative images of immunolabelled integrin α11 (green) and the lysosomal marker Lamp1 (magenta) in control-treated (ctrl) or Bafilomycin A1-treated (100 nM, 6h) wild-type and *Sharpin^cpdm^* MSFs plated on collagen. Region of interest (yellow) shows examples of co-localization (white spots, highlighted with red arrow). (**H**) Quantification of relative (to wt control) co-localization of integrin α11 and Lamp1 in control-treated or Bafilomycin A1-treated wild-type and *Sharpin^cpdm^* MSFs plated on collagen, n= 76, 102, 89, 125 cells (from left to right) pooled from 3 independent experiments, line under p-value indicates which samples are compared with each other. Mean ± SEM in all graphs. (A) Wilcoxon matched-pairs signed rank test. (C, F, H) Mann-Whitney *U*-test. Scale bars: 20 μm.

As inactive and active integrins are trafficked and degraded differently (Arjonen et al., 2012; Rainero et al., 2015), we speculated that the increased relative activity of integrin β1 could lead to increased targeting of integrin α11β1 to lysosomal degradation in *Sharpin^cpdm^* MSFs. This hypothesis was supported by the fact that SHARPIN-deficient MSFs displayed more integrin α11 co-localization with Lamp1 (lysosomal marker) than wild-type MSFs (Fig 4G, H). Furthermore, we observed accumulation of integrin α11 to the Lamp1-positive compartment when MSFs were treated with the lysosomal degradation disruptor Bafilomycin A1 (Fig. 4G, H). These data suggest that a subset of integrin α11 traffics to lysosomal degradation in MSFs and that this is enhanced in SHARPIN deficient cells.

To investigate whether the reduced surface expression of integrin α1 and α11 in *Sharpin^cpdm^* MSFs could be responsible for the impaired capability to spread on a compliant collagen I-coated surface, we analyzed cell spreading following siRNA-mediated downregulation of integrin α1 or integrin α11 (Fig. 5A), confirmed by qPCR (Fig. S3E). In wild-type MSFs downregulation of integrin α11, but not integrin α1, led to significantly reduced cell spreading on 2 kPa collagen I-coated hydrogels that resembled the phenotype of *Sharpin^cpdm^* MSFs (Fig. 1B). In contrast, no significant differences in cell area were observed when the endogenously lower integrin α1 and integrin α11 levels were silenced in *Sharpin^cpdm^* MSFs (Fig. 5A). These data indicate that appropriate levels of integrin α11, but not integrin α1, are important for regulating the spreading of MSFs on soft collagen I-coated substrates. This could be linked to the fact that integrin α11 mediates strong binding to and contraction of fibrillar collagen I (Tiger et al., 2001) whereas integrin α1 prefers non-fibrillar collagen IV and has lower binding affinity to fibrillar collagens (types I, II and III) (Tulla et al., 2001). However, this remains to be investigated.

**Figure 5.**
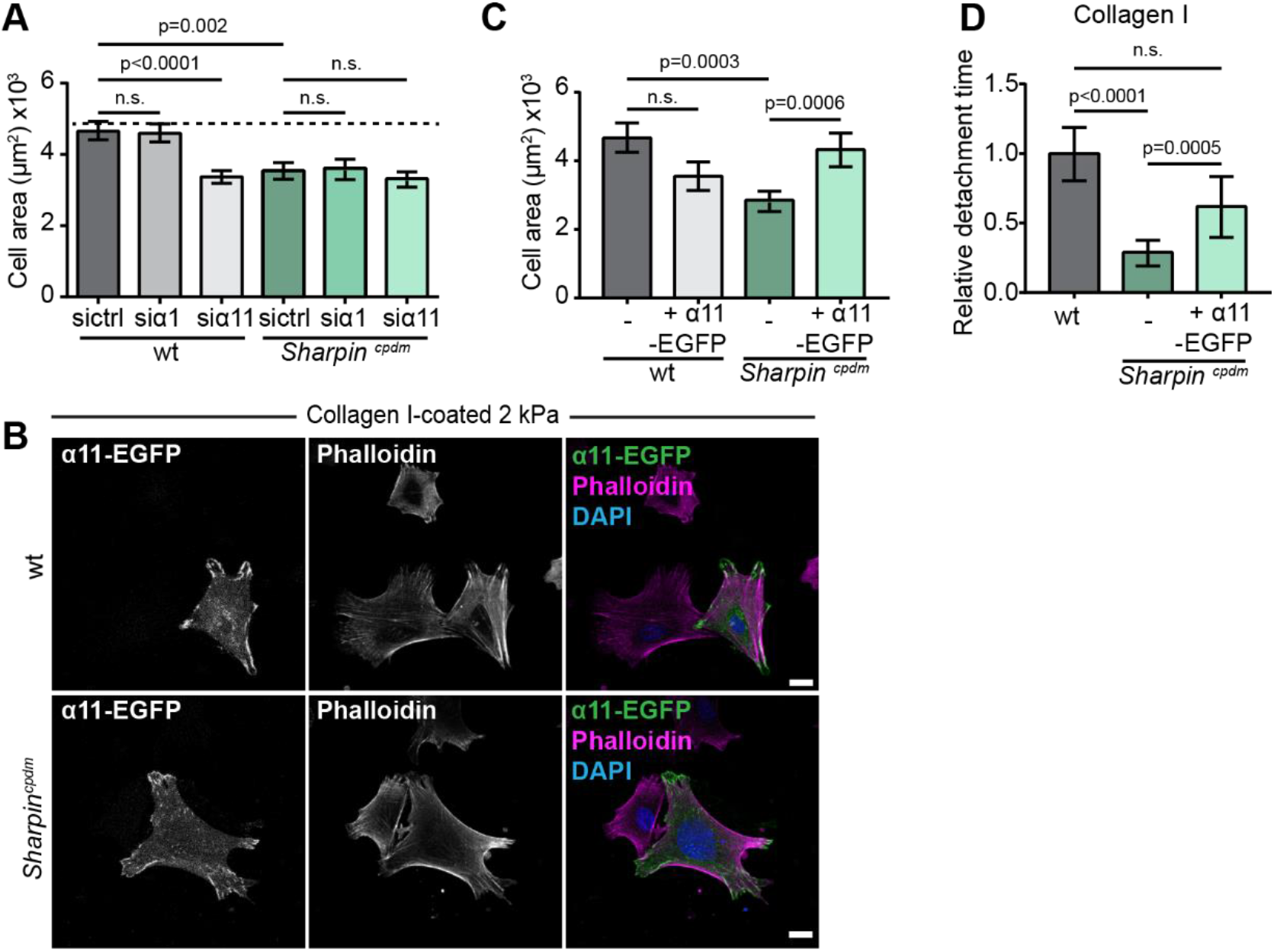
Integrin α11β1 regulates the spreading of MSFs on soft matrices. (**A**) Quantification of the cell area in wild-type and *Sharpin^cpdm^* MSFs silenced with control, integrin α1 or integrin α11 targeting siRNA and plated on 2 kPa collagen I-coated PAA hydrogels; n=94, 91, 93, 82, 88, 90 cells (from left to right) from three independent experiments. (**B**) Representative images of integrin α11-EGFP transfected wild-type and *Sharpin^cpdm^* MSFs plated on 2 kPa collagen I-coated PAA hydrogels, and co-labelled for F-actin (magenta) and nuclei (blue). (**C**) Quantification of cell area in non-transfected and integrin α11-EGFP transfected wild-type and *Sharpin^cpdm^* MSFs plated on 2 kPa collagen I-coated PAA hydrogels. Data are pooled from two independent experiments, n=37, 31, 46, 48 cells (from left to right). (**D**) Quantification of detachment time of wild-type, *Sharpin^cpdm^* and integrin α11-EGFP transfected *Sharpin^cpdm^* MSFs from collagen I-coated magnetic beads. Data are pooled from three independent experiments, n=91, 101, 46 cells (from left to right). Mean ± SEM in all graphs. (A) unpaired t-test, (C, D) Mann-Whitney *U*-test. Scale bars: 20 μm.

To verify that integrin α11 levels are critical for collagen I interaction of MSFs, we performed rescue experiments with ectopically expressed EGFP-tagged human integrin α11. Reintroduction of integrin α11 in *Sharpin^cpdm^* MSFs on collagen I-coated 2 kPa hydrogels reversed their defective spreading (Fig. 5B, C). In contrast, overexpression of integrin α11-EGFP in wild-type MSFs modestly, albeit, non-significantly decreased cell spreading (Fig. 5B, C). In addition, reintroduction of integrin α11 by ectopic expression partially rescued the ligand detachment time in *Sharpin^cpdm^* MSFs (Fig. 5D). Taken together, these results demonstrate that integrin α11β1 is essential in the integrin-collagen I binding dynamics and spreading of MSFs on soft collagen I-coated substrates, and that the impaired mechanotransduction of SHARPIN-deficient MSFs is coupled to reduced integrin α11β1 protein expression level.

## Discussion

Here we have investigated the mechanotransduction of mammary gland stromal fibroblasts to gain insight into the biologically essential role of these cells in sculpting the mammary gland stromal architecture *in vivo* (Peuhu, Kaukonen et al., 2017). Taking this experimentally challenging primary cell model (compared to the immortalized cell lines studied previously) has provided two interesting and unexpected observations related to tissue specific characteristics of mechanotransduction. First, our data comparing wild-type and SHARPIN-deficient cells provide a striking example of how, even if major mechanical regulators such as myosin contractility are not affected, merely changing integrin properties under force can dramatically affect the cell’s mechanoresponse. Second, the fact that wild-type MSFs exhibit mature, vinculin-associated FA irrespective of matrix stiffness allowed us to decouple traction force generation from adhesion maturation, leading to the fundamental clutch model prediction of a biphasic traction-stiffness relationship that is otherwise very elusive to observe (Elosegui-Artola et al., 2018).

The differential regulation of collagen- and fibronectin-binding integrins, and the mechanobiological implications of these differences remain poorly understood. Here, we have investigated the consequences of SHARPIN-deficiency on cell mechanosensing, integrin ligand-binding dynamics, and traction force generation, and conducted computational modelling of these events. Our data demonstrate that in the absence of the integrin activity regulator SHARPIN, the protein levels of the collagen-binding integrin α11β1 are downregulated. This is likely to be the main reason for the significantly increased unbinding of SHARPIN-deficient MSFs from collagen-coated beads under force. As the remaining integrin α11β1 is more likely to be in the primed active conformation, due to the increase in relative integrin β1 activity, this could account for the small increase of integrin recruitment to collagen. These faster cell-collagen binding dynamics are in line with the previously reported rapid adhesion turnover of SHARPIN-deficient MSFs (Peuhu et al. 2017). Importantly, the molecular clutch model was able to predict the absence of traction peak and increased actin flow rate at low rigidities based on increased clutch binding and unbinding rates to collagen, measured by magnetic tweezers in SHARPIN-deficient cells. Of note, whereas the response to force of integrin-collagen bonds is not known, our model assumed a catch bond behaviour. This should however not affect our conclusions, since the key aspects of the molecular clutch model found here (biphasic behaviour, plus the role of fold changes in binding and unbinding rates) are general to both catch and slip bonds (Bangasser et al., 2013).

Our model failed, however, to predict the final increase in force transmission in wild-type cells with increasing stiffness. While the details of this regime warrants further investigation, we note that in conditions with very low actin flow rates, mature adhesions and very stable actin fibers, cytoskeletal reorganization events other than fast actin flow could be determinant of the cellular force transmission (Oakes et al., 2012). In all, we acknowledge that the model prediction corresponds to the experimental data only in the soft stiffness range, possibly because not all cellular parameters have been, and can be, taken into account by the molecular clutch model.

Although SHARPIN regulates the activity of both collagen and fibronectin-binding integrins (Rantala et al., 2011), the low levels of fibrillar collagen binding integrin α11 in SHARPIN-deficient MSFs results in different mechanosensitive responses of these cells on collagen and fibronectin. This is in line with the established role of integrin α11β1 (Lehnert et al., 1999), in the regulation of collagen contractility (Popova et al., 2007; Tiger et al., 2001) and in collagen-linked disease conditions, such as fibrosis, cancer invasion and particularly in cancer-associated fibroblasts (Bansal et al., 2017; Navab et al., 2016). The regulatory pathways modulating the activity of collagen-binding integrins may be distinct from other integrin heterodimers. The vast majority of studies investigating integrin activation are based on the platelet-specific integrin αIIbβ3 and the fibronectin receptors integrin α5β1 and αvβ3, which are primarily regulated by inside-out and outside-in signaling. In turn, only a few studies have addressed activity regulation in the context of collagen-binding integrins. Heterodimerization of α1β1 and α2β1 integrins has been postulated to have a key role in their activity regulation based on the lower affinity of integrin α1 and α2 to integrin β1 (Lu et al., 2016). Thus, regulation of the expression levels of collagen-binding integrins may be particularly important for their ligand binding dynamics.

SHARPIN is a multifunctional adapter protein that has been implicated in a number of other signaling pathways, including inhibition of integrin activity (Kasirer-Friede et al., 2019; Peuhu, Kaukonen et al., 2017; Peuhu, Salomaa et al., 2017; Pouwels et al., 2013; Rantala et al., 2011). SHARPIN promotes canonical Nuclear factor (NF)-κB activation (Gerlach et al., 2011; Tokunaga et al., 2009) and other inflammatory signaling cascades as part of the linear ubiquitin assembly complex (LUBAC) (Chattopadhyay et al., 2016; Dubois et al., 2014; Rodgers et al., 2014; Zak et al., 2011). Furthermore, SHARPIN regulates the functions of the Arp2/3 protein complex (Khan et al., 2017), T cell receptor (Park et al., 2016), and caspase 1 (Nastase et al., 2016) in a LUBAC-independent manner, and interacts with PTEN (He et al., 2010), SHANK proteins (Lim et al., 2001), and EYA transcription factors (Landgraf et al., 2010). In this study, reduced integrin α11 protein levels were directly linked to mechanobiological phenotypes in SHARPIN-deficient MSFs, while integrin expression at the transcriptional level remained comparable to wild-type MSFs. Since NF-κB is a transcription factor that functions predominantly through regulation of gene expression, our data imply that Nf-kB might not be the primary mechanism involved in the regulation of integrin α11 in SHARPIN-deficient MSFs. Currently, we lack the detailed mechanisms accounting for the reduced integrin α11 protein levels in SHARPIN-deficient MSFs and in wild-type MSFs upon SHARPIN silencing. However, it is likely that increased lysosomal trafficking of integrin α11, in the absence of SHARPIN, is contributing to elevated integrin degradation in these cells. This would be in line with previous studies linking integrin activation to reduced receptor recycling rates and increased lysosomal degradation (De Franceschi et al., 2015).

Together, our findings demonstrate how altered integrin activity in SHARPIN-deficient primary MSFs results in deregulated cell spreading and traction force generation in response to substrate ligand composition and stiffness. The central role for integrin α11, uncovered here, in regulating mechanotransduction on collagen may also be essential to the pathological behavior of fibroblasts in cancerous or fibrotic tissues. As both SHARPIN and integrin α11β1 are significant regulators of cancer tumorigenesis and dissemination, as well as fibroblast contractility and collagen remodeling (He et al., 2010; Navab et al., 2016; Peuhu, Kaukonen et al., 2017; Tamiya et al., 2018; Zhu et al., 2007), increased understanding of their functional interplay is of wide interest. Finally, our finding that the mechanical output of fibroblasts can be strongly influenced by a single parameter of the molecular clutch, the integrin binding dynamics, highlights the tuneability of mechanotransduction, and its ability to trigger specific outputs in response to both internal and external parameters.

## Materials and methods

### Animals

The C57BL/KaLawRij-*Sharpin^cpdm^*/RijSunJ mouse strain (Stock No: 007599) with a spontaneous chronic proliferative dermatitis *(cpdm)* mutation in *Sharpin* gene leading to the complete loss of SHARPIN protein (HogenEsch et al., 1993; Seymour et al., 2007) was acquired from The Jackson Laboratory (Bar Harbor, ME). The colony was maintained and genotyped as previously described (Peuhu, Kaukonen et al., 2017). Six to seven week-old, female *Sharpin^cpdm^ (Sharpin^cpdm/cdpm^)* mice and littermate wild-type mice *(Sharpin^+/+^* or *Sharpin^+/cpdm^)* were used for MSFs isolation. Mice were housed in standard conditions (12-h light/dark cycle) with food and water available *ad libitum.* The viability, clinical signs and behaviour of the mice were monitored daily. For euthanasia, cervical dislocation was used in conjunction with CO_2_. All animal experiments were ethically assessed and authorised by the National Animal Experiment Board and in accordance with The Finnish Act on Animal Experimentation (Animal licence number ESAVI-9339-04.10.07-2016)

### Isolation of primary cells

Isolation of MSFs was performed as previously described (Peuhu, Kaukonen et al., 2017). Briefly, mouse mammary glands were dissected after sacrifice, lymph nodes removed and the tissue moved to ice cold PBS. The tissue was first mechanically disrupted by mincing with scalpel followed by 2-3 h enzymatic digestion (1% L-glutamine, insulin 5 μg/ml, gentamycin 50 μg/ml, 5% FCS and 2 mg/ml collagenase in DMEM/F-12), and DNAse I treatment (20 U/ml). Fibroblasts were isolated by repeated pulse centrifugation (1500 rpm) and collecting each time the supernatant that contained the stromal cells. After the last pulse centrifugation, the collected supernatant was pelleted and resuspended in growth medium (1% L-glutamine, 1% penicillin/streptomycin, 5% FCS in DMEM/F-12) and the cells plated for culture. Medium was replaced the next day to remove non-adherent and dead cells. Throughout the study, cells were grown for a maximum of 5 passages before being discarded and experiments were performed on multiple cell isolations each of which originated from 2-4 animals.

### Cell culture

The talin1^−/−^ MEF cell line was described previously (Elosegui-Artola et al., 2016) and was cultured in DMEM containing 15 % FCS.

### Transient transfection and gene silencing

Plasmids [mEmerald-Lifeact-7 (addgene, #54148) and hITGA11-GFP (pBJ1 human integrin alpha 11-EGFP; Erusappan, P., et al. Integrin α11 cytoplasmic tail is required for FAK activation to initiate 3D cell invasion and ERK-mediated cell proliferation. Manuscript in preparation)] were transfected using DNA-In reagent (GST-2131, MTI Global Stem) or Lipofectamine 3000 (L3000075, ThermoFisher) according to the manufacturer’s instructions. Briefly a transfection mixture of DNA-In reagent and plasmid was prepared in 1:3 ratio (μg DNA: μl DNA:In) in 250 μl optimum and incubated for 15 min before transfection. The transfection mixture was added to adhered cells grown on a 6-well plate in 2.5 ml fresh media and cells incubated for 24 h. For transfection with Lipofectamine 3000 reagent a mixture of ratio 4:4:2.5 (μl Lipofectamine 3000 reagent: μl P3000TM Enhancer reagent: μg DNA) was prepared in 500 μl optimem and incubated for 15 minutes before adding to adhered cells grown on a 6-well plate in 2 ml fresh antibiotic free media and cells incubated for 24 h.

ON-TARGETplus Mouse Itga1 (109700; 5’-CUUUAAUGACGUCGUGAUU-3’, 5’-GCCUAUGACUGGAACGGAA-3’, 5’-CCACAAUUGACAUCGACAA-3’, and5’-AGGGCAAGGUGUACGUGUA-3’) and Mouse Itga11 (319480; 5’-AUGGAUGAGAGGCGGUAUA-3’, 5’-UCAGAAGACAGGAGACGUA-3’, 5’-GCAUCGAGUGUGUGAACGA-3’, and 5’-CCAGCGACCCUGACGACAA-3’) siRNA –SMARTpools were ordered from Dharmacon, and SHARPIN siRNA (5’-GCUAGUAAUUAAAGACACAd(TT)-3’) and the scramble Allstars negative control siRNA were ordered from QIAGEN. Gene silencing was performed using siRNA oligonucleotides and Lipofectamine RNAiMax reagent (13778150, Thermo Fisher Scientific) according to the manufacturer’s protocol. Briefly a mixture in 4:3 ratio (μl RNAiMax: μl siRNA) was prepared in 200 μl optimum and incubated for 20 min before adding to adhered cells grown on a 12-well plate in 400 μl fresh media and cells incubated for 48 h.

### Preparation of polyacryl-amide hydrogels

35 mm glass bottom dishes with 14 mm bottom wells (Cellvis, catalog number D35-14-1-N) were treated with 1 ml Bind silane solution (7.14% Plus One Bind Silane (Sigma, GE17-1330-01), 7.14% acetic acid in 96 % ethanol) for 30 min, washed twice with 96 % ethanol and left to dry completely. A hydrogel mixture containing 7.5-18 % acrylamide solution (Sigma) and 0.06-0.4 % bis acryl amide solution (Sigma), diluted in PBS up to a final volume of 500 μl was prepared to obtain hydrogels ranging in stiffness from 0.75-13 kPa. The polymerization of the mixture was initialized by adding 5 μl 10 % ammonium persulfate (BioRad) and 1 μl N,N,N’,N’-Tetramethylethylenediamine (Sigma). The hydrogel solution was quickly vortexed, 11.7 μl was added on top of the glass bottom dish, and a 13 mm glass coverslip carefully placed on top of the drop and the gel was let to polymerize for 1 h RT incubation. After polymerization, the coverslip was carefully removed and the gel incubated with PBS to prevent drying. The stiffness of the hydrogels was confirmed by atomic force microscopy as previously described (Närvä et al., 2017).

### Gel surface activation and coating

Gels were incubated for 30 min on slow agitation with 500 μl Sulfo-SANPAH activation solution [0.2 mg/ml Sulfo-SANPAH (Sigma, catalog number 803332), 2 mg/ml N-(3-Dimethylaminopropyl)-N’-ethylcarbodiimide hydrochloride (Sigma, catalog number 03450) in 50 mM Hepes] followed by 10 min UV-light activation. After the activation gels were washed three times with PBS and coated with either fibronectin (Merck-Millipore, catalog number 341631) (20 μg/ml) or collagen type I (Merck-Millipore, catalog number 08-115) (20 μg/ml).

### Immunofluorescence

Cells were fixed and permeabilized with 0.1 % triton in 4 % PFA for 8 minutes followed by 10 min fixation with 4 % PFA. When Bafilomycin A1 was used, cells were treated with 100 nM Bafilomycin A1 (Merck-Millipore, catalog number 196000) solubilized in DMSO for 6h prior to fixation. For antibody staining against integrin α11 samples were fixed with methanol for 10 min −20 C, followed by 10 min 0.1 % triton permeabilization in RT. To block unspecific binding of antibodies cells were incubated in 10 % horse serum (HRS) for 1 h in RT. Primary and secondary antibodies were diluted in 10 % HRS and incubated for 1h in RT. Primary antibodies used: mouse (ms) anti-vinculin (V9131, Sigma-Aldrich, 1:500), rat anti-LAMP1 ([1D4B], ab25245, Abcam, 1:100), ms anti-YAP (sc-101199, Santa Cruz, 1:100), rabbit (rbt) anti-mouse integrin α11 (provided by Donald Gullberg, 1:200) and rat anti-integrin β1, clone MB1.2 (LV1766450, Millipore, 1:100). Secondary antibodies used: Alexa Fluor 488 anti-mouse (A-21202, Thermo Fisher, 1:400), Alexa Fluor 488 anti-rabbit (A-21206, Thermo Fisher, 1:400), Alexa Fluor 647 anti-rat (A-21247, Thermo Fisher 1:400) and Alexa Fluor 594 anti-rat (A21209, Thermo Fisher 1:400). F-actin was stained with Phalloidin–Atto 647N (65906, Sigma, 1:400), incubated together with secondary antibodies and nuclei with DAPI (D1306, Life Technologies 1:3000) for 10 min RT after secondary antibody incubation.

### Quantification of immunofluorescence imaging

Quantification of the cell area and the number, average size, and average length (Feret diameter ie. the longest distance between any two points along the object boundary, also known as maximum caliper) of adhesions per cell was based on vinculin immunofluorescence labeling. Quantification of the nuclear:cytoplasmic ratio of YAP was performed with CellProfiler from maximum intensity projection (MIP) images. Nuclei were segmented based on the DAPI staining, mean intensity was quantified for the nuclei and the surrounding cytoplasms. Quantification of the co-localization of integrin α11 and Lamp1 was performed with the ImageJ plugin ComDet, `https://github.com/ekatrukha/ComDet (Fréal et al., 2019), which ignores inhomogeneous cytoplasmic background. We detected integrin α11 spots (pixel size 4) that colocalized with Lamp1 spots based on the distance between them (pixel distance 3). Then the percentage of integrin α11 spots that co-localized with Lamp1 spots was calculated as the number of colocalizing integrin α11 spots of the total number of integrin α11 spots.

### Confocal imaging

Samples were imaged using 3i (Intelligent Imaging Innovations, 3i Inc) Marianas Spinning disk confocal microscope with a Yokogawa CSU-W1 scanner and Hamamatsu sCMOS Orca Flash 4.0 camera (Hamamatsu Photonics K.K.) using 40x/1.1 water objective), LSM 880 Airyscan laserscanning confocal microscope (Zeiss) using 20x/0.8 objective, 63x/ 1.4 oil objective or LSM 880 Airyscan LD LCI confocal microscope (Zeiss) using Plan-Apochromat 40x/1.2 water objective and Airyscan detector.

### Flow cytometry

MSFs were analysed by flow cytometry as previously described (Peuhu, Kaukonen et al., 2017). Briefly, cells were detached, placed on ice, fixed with 4% PFA in PBS (10 min RT) and resuspended in PBS. Cell surface integrins were labelled with fluorochrome-conjugated antibodies [Alexa Fluor 488 anti-integrin β1 (Clone: HMβ1-1, Biolegend), APC anti-integrin α1 (Clone: HMα1, Biolegend), or Alexa Fluor 488 anti-integrin α5 (Clone: 5H10-27, Biolegend)] diluted in 100 μL Tyrode’s buffer (10 mM HEPES-NaOH at pH 7.5, 137 mM NaCl, 2.68 mM KCl, 1.7 mM MgCl2, 11.9 mM NaHCO3, 5 mM glucose, 0.1% BSA) according to manufacturer’s instrctions for 30 min at RT. Alternatively, cells were first labelled with primary antibodies [integrin α11 (AF6498, R&D), or active integrin β1 (clone 9EG7, BD Pharmingen)] followed by secondary antibodies [Alexa Fluor 647 anti-sheep (ab150179, Abcam) or Alexa Fluor 488 anti-rat (Molecular Probes)] both dilutedin Tyrode’s buffer. After washes, the samples were analysed using BD LSRFortessa flow cytometer. Live cells were gated from FSC-A/SSC-A dot blot and analysed for geometric mean or median fluorescence intensity. The expression of active integrin β1 was normalized to the total surface expression of integrin β1.

### Western blotting

Because of the small hydrogel area (13 mm) on our in-house hydrogels we used the commercial hydrogels Softwell^®^ 6, Easy Coat™ (SW6-EC-0.5, SW6-EC-2 EA, SW6-EC-4 EA and SW6-EC-50 EA, Matrigen) for immunoblotting samples collected from hydrogels.

Protein extracts were prepared by lysing the cells with hot TX lysis buffer [50 mM Tris–HCl pH 7.5, 150 mM NaCl, 5% glycerol, 1% SDS, 0.5% Triton X-100, Complete protease inhibitor, PhosSTOP (Roche)]. Samples were sonicated with BioRuptor and protein concentration was measured by BioRad to assure equal protein loading. The protein extract was first separated by loading equal amounts of protein on 4-20 % Mini-PROTEAN^®^ TGX™ Gel SDS–PAGE gradient gels (456-1096, Biorad) and then transferred to the nitrocellulose membrane with Trans-Blot Turbo Transfer Pack (170-4159, Biorad). The membrane was blocked for 1 h in RT with 5% milk in Tris Buffered Saline and 0.1% Tween 20 (TBST) solution before antibody incubation. Primary and secondary antibodies were incubated for a minimum of 1 h. Membranes were scanned and results analyzed with the Odyssey infrared system (LICOR Biosciences). Primary antibodies used for western blotting: Rabbit anti-Phospho-Myosin Light Chain 2 (Thr18/Ser19) (3674, Cell Signaling Technology), rabbit anti-Myosin Light Chain 2 (3672, Cell Signaling Technology), mouse anti-glyceraldehyde-3-phosphate dehydrogenase (GAPDH) (5G4-MAb: 6C5, HyTest), rabbit anti-SHARPIN (14626-1-AP, Proteintech) and rabbit anti-Integrin α11 (provided by D. Gullberg). Secondary antibodies used for western blotting: IRDye^®^ 800CW Donkey anti-Mouse IgG, IRDye^®^ 800CW Donkey anti-Rabbit IgG, IRDye^®^ 680LT Donkey anti-Mouse IgG and IRDye^®^ 680LT Donkey anti-Rabbit IgG, diluted 1:10000 in odyssey blocking buffer (LI-COR). Protein levels were determined by comparing the signal intensity value of the protein of interest relative to the signal intensity value of the corresponding loading control. The relative intensity value was then divided by the sum of the relative intensity values in each independent experiment.

### Actin flow

EGFP-LifeAct transfected cells were plated on 2 kPa collagen coated hydrogels and allowed to spread for 45-105 min before image acquisition. Imaging was performed with a Carl Zeiss LSM 880 Airyscan microscope using a 63x/1.4 oil immersion objective and its airyscan detector array. Images were acquired every second for 125 seconds and actin flow was measured based on the slope from one kymograph per cell drawn along actin fibers close to the cell periphery at the leading edge.

### Traction force microscopy

For traction force microscopy experiments, cells were seeded 4h before performing experiments on polyacrylamide hydrogels of different stiffness prepared as previously described (Elosegui-Artola et al., 2016). Then, simultaneous images were acquired of single cells (phase contrast) and of fluorescent 200 nm beads embedded in gels. Images were acquired with a Nikon Ti Epifluorescence microscope with a 40x objective (N.A. 0.6). Afterwards, cells were trypsinized, and images of bead positions in the gel in the relaxed state were acquired. By comparing bead positions in the deformed versus relaxed positions, a map of gel displacement caused by cells was measured using a custom particle-imaging-velocimetry software (Bazellières et al., 2015). Then, assuming that the displacements were caused by forces exerted by cells on the cell-gel contact area, forces were measured using a previously described Fourier transform algorithm (Butler et al., 2002; Oria et al., 2017). The average forces per unit area of each cell was then measured.

### Magnetic tweezers and bead recruitment experiments

Magnetic tweezers experiments were performed as previously described (Elosegui-Artola et al., 2014; González-Tarragó et al., 2017; Roca-Cusachs et al., 2009). Briefly, 3 μm carboxylated magnetic beads (Invitrogen) were coated with a mixture of biotinylated BSA and either biotinylated fibronectin or collagen I fragment (10:1 ratio). This fragment was pentameric FN7-10 for fibronectin (Coussen et al., 2002) and the GFOGER peptide for collagen I (Emsley et al., 2000). Two hours after seeding cells on glass coverslips, coated magnetic beads were deposited on top of the coverslips and allowed to attach to cells. Then, magnetic beads on the lamellipodia of single cells were pulled with a 1 nN pulsatory force (1 Hz), and the time required to detach beads was measured. To quantify the recruitment of integrin β1, 3 μm carboxylated silica beads (Kisker Biotech) were used and coated as described above. Instead of pulling the beads, cells with silica beads were fixed and stained for integrin β1 (Abcam, 12G10). The average intensity of both beads and surrounding areas was quantified, and the difference between those values was taken as the integrin recruitment measure.

### Mathematical modelling

Modelling was carried out using the molecular clutch model previously described in detail (Elosegui-Artola et al., 2014). Briefly, the model considers a given number of myosin motors, pulling on an actin bundle. The actin bundle can bind to a set of collagen ligands through molecular clutches that represent adaptor proteins and integrins. In turn, collagen ligands are connected to the substrate through a spring constant representing substrate stiffness. Molecular clutches bind to the collagen ligands with an effective binding rate, and unbind with an unbinding rate that depends on force as a catch bond. The clutches transmit forces to the substrate only when they are bound, and therefore overall force transmission critically depends on binding dynamics. In simulations, all parameters remain constant except the substrate spring constant (which increases with stiffness), and the number of myosin motors pulling on actin, which increases with stiffness following the results in Figure 4D, E. This increase is the same for both wild-type and *Sharpin^cpdm^* cells. The only parameters that are different in the model between both conditions (wild-type and *Sharpin^cpdm^)* are the unbinding and binding rates, which increase by the same proportion (8-fold) in *Sharpin^cpdm^* cells. We note that in contrast to our previous work (Elosegui-Artola et al., 2014; Elosegui-Artola et al., 2016; Oria et al., 2017), in this case we do not introduce reinforcement in the model, i.e. force-dependent recruitment of additional integrins. This is because focal adhesion size and distribution is largely independent of both stiffness and SHARPIN-deficiency. We also note that for unbinding rate, we took the catch bond force dependence reported in (Kong et al., 2009) for α5β1 integrins. This dependency is likely to change for collagen-binding integrins, for which unfortunately there are to our knowledge no reported measurements of systematic force-lifetime measurements at the single molecule level. However, whereas specific levels of force depend on this dependence, overall trends with stiffness, and the relative differences if overall unbinding rates are altered (as occurs upon SHARPIN depletion) are maintained regardless of the specific force dependence assumed. See Table S1 for a list of parameters.

### Statistical analysis

GraphPad software was used for all statistical analyses. Student’s *t*-test (unpaired, two-tailed) was used when normality could be confirmed by D’Agostino& Pearson omnibus normality test. Nonparametric Mann–Whitney *U*-test was used when two non-normally distributed groups were compared or when normality could not be tested [due to a too small data set (*n*< 8)]. Wilcoxon matched-pairs signed rank test was used if samples with unequal variance were compared. Fisher’s exact test was used for the analysis of contingency tables. Data are presented in column graphs with mean ± standard error of mean (SEM) and *P*-values. Individual data points per condition are shown when *n* ≤ 15, and *n*-numbers are indicated in figure legends.

## Author contributions

PRC, EP and JI contributed to the conception and design of the study. ML, AEA, MG, JZK and EP designed, conducted and analysed *in vitro* experiments. AEA and JZK performed traction force microscopy. ML and AEA conducted the magnetic tweezer experiments. CG analysed the TFM data and confirmed gel stiffness with AFM. ML, EP and JI wrote the manuscript, and AEA, PRC, and DG edited the manuscript. DG, PRC, EP and JI supervised the research. Funding acquisition AEA and JI.

## Acknowledgments

We thank J. Siivonen, and P. Laasola for technical assistance; P. Erusappan for integrin α11 antibody protocols; A. Isomursu and the rest of the Ivaska Lab for insightful comments; H. Hamidi for editing of the manuscript; and X. Trepat for thought provoking discussions that inspired this study. Turku Bioscience Cell Imaging Core, Turku Center for Disease Modeling and Biocenter Finland are acknowledged for services, instrumentation and expertise. This study has been supported by the Academy of Finland (Grant 312517), ERC Consolidator Grant (615258), the Sigrid Juselius Foundation, and the Finnish Cancer Organization. ML has been supported by Turku Doctoral Program of Molecular Medicine, Instrumentarium Science Foundation and The Swedish Cultural Foundation in Finland, and EP by Academy of Finland and Finnish Cultural foundation.

---

**Figure S1.**
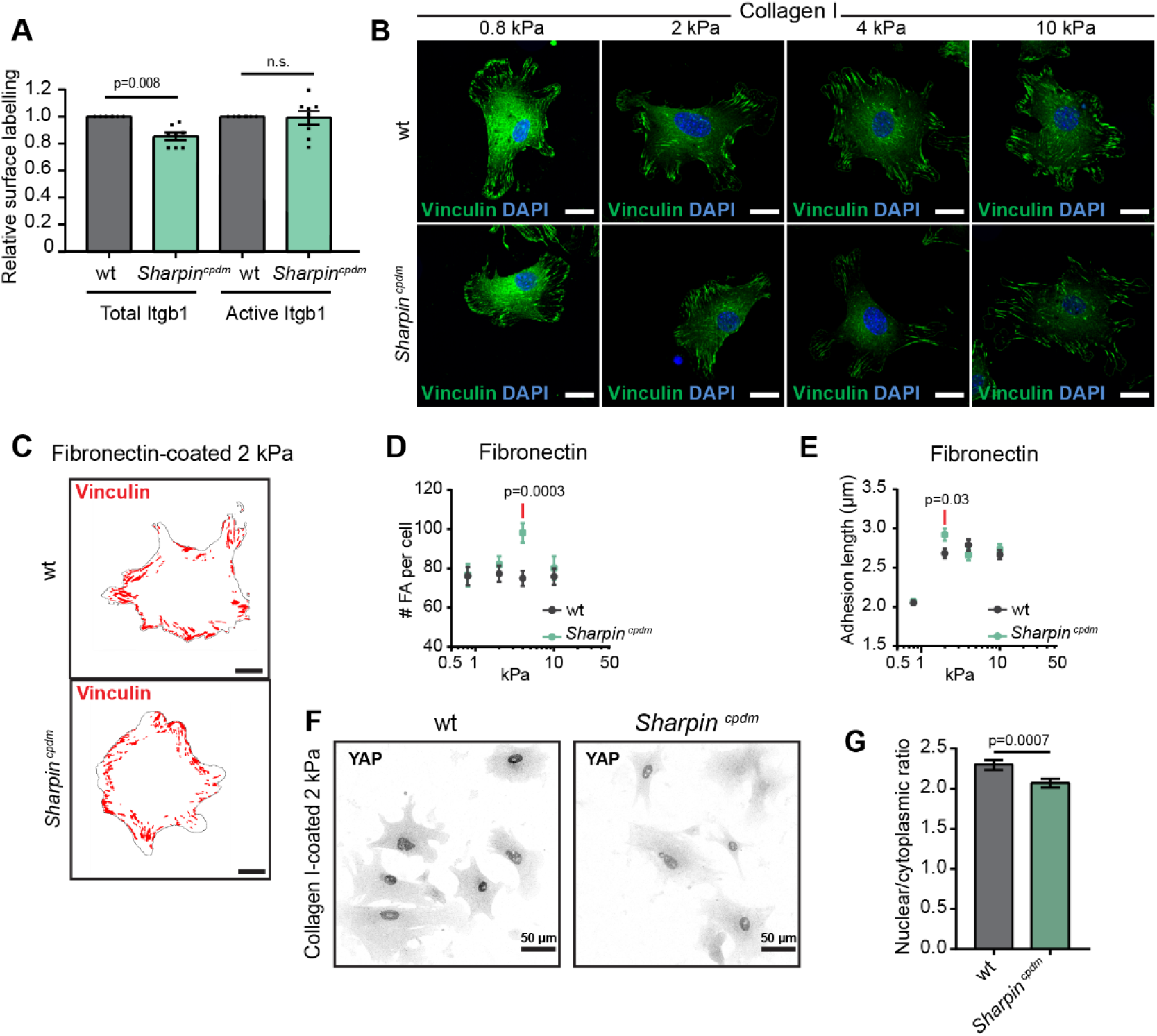
(**A**) Measurement of total integrin β1 (clone, HMβ1-1) and active integrin β1 (clone, 9EG7) cell-surface levels (n=7 independent experiments) in *Sharpin^cpdm^* relative to wild-type MSFs by flow cytometry. (**B**) Representative images of vinculin-containing FAs (green) in wild-type and *Sharpin^cpdm^* MSFs plated for 3-4 h on 0.8, 2, 4 and 13 kPa collagen I-coated PAA hydrogels, nuclei (blue) were colabelled. (**C**) Representative output images of FA analysis for individual cells plated on 2 kPa fibronectin-coated PAA hydrogels (FA, red; cell borders, black). (**D, E**) Quantification of the number of FA per cell (D) and the length of FA (E) in wild-type compared to *Sharpin^cpdm^* MSFs plated on fibronectin-coated 0.8, 2, 4 and 13 kPa PAA hydrogels. Data are pooled from three independent experiments, n_wt_= 89, 95, 103, 99 and n_Sharpincpdm_= 66, 79, 82, 82 cells (# FA per cell, from left to right) and n_wt_= 89, 95, 103, 99 and n_Sharpincpdm_= 66, 78, 82, 82 cells (Adhesion length, from left to right). (**F**) Representative images of YAP immunolabeling in wild-type and *Sharpin^cpdm^* MSFs plated on collagen I-coated 2 kPa PAA hydrogels and (**G**) quantification of nuclear to cytoplasmic localization ratio of YAP. Data are pooled from three independent experiments, n_wt_=156 and n_Sharpincpdm_=240 cells. Mean ± SEM in all graphs. Mann–Whitney *U*-test, red lines above and black lines below p-values indicate the data points compared. (B, C) Scale bars: 20 μm.

**Figure S2.**
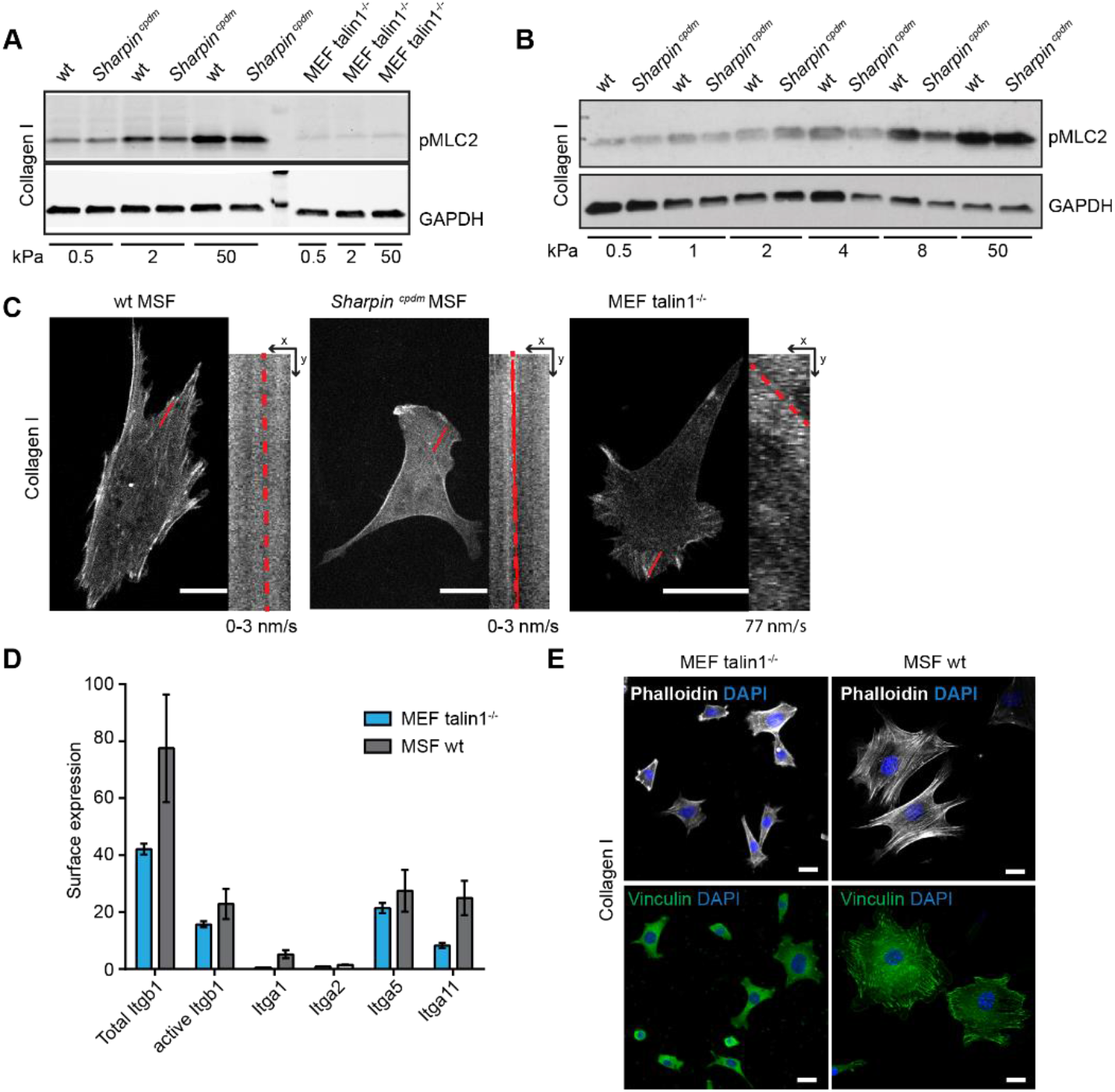
(**A**) Representative Western blot analysis of pMLC2 and GAPDH protein expression in wild-type and *Sharpin^cpdm^* MSFs or in Talin1−/− MEFs plated on PAA hydrogels with the indicated stiffness. (**B**) Representative Western blot analysis of pMLC2 and GAPDH protein expression in control and *Sharpin^cpdm^* MSFs plated on PAA hydrogels with the indicated stiffness. (**C**) Representative images of Lifeact-GFP transfected wild-type and *Sharpin^cpdm^* MSFs or Talin1−/− MEFs plated for 2-3h on 2 kPa collagen-coated PAA hydrogels. Insets are kymographs showing actin retrograde flow along the red line (time=180s, imaged every second). The slope of the line was used to calculate the actin retrograde flow rate. (**D**) Measurement of cell-surface expression of integrin β1, active integrin β1, integrin α1, α5 and α11 in Talin1−/− MEFs and wild-type MSFs, (normalized to secondary control staining), n=4 independent experiments. (**E**) Representative images of Talin1−/− MEFs and wild-type MSFs plated for 3-4 h on 2 kPa collagen I-coated PAA hydrogels and labelled for F-actin (white, upper panel) or vinculin (green, lower panel), nuclei (blue) was co-labelled. Scale bars: 20 μm.

**Figure S3.**
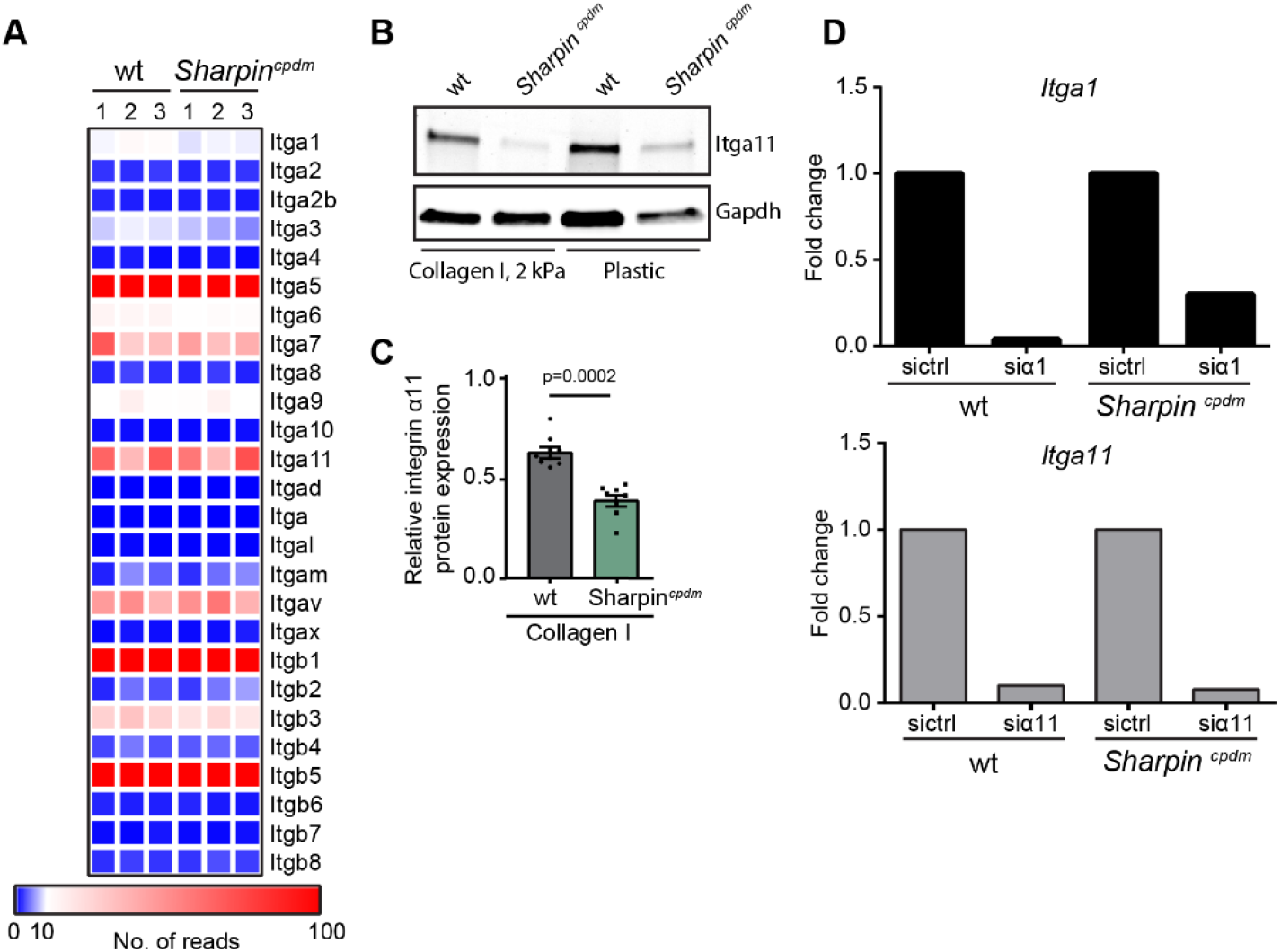
(**A**) Heat map of integrin mRNA expression in wild-type and *Sharpin^cpdm^* MSFs detected by RNA sequencing (Peuhu, Kaukonen et al., 2017). The normalized number of mRNA reads from three independent cell isolation replicates is shown. Colour scale represents a range from 0-100 reads. (**B**) Western blot analysis of integrin α11 and GAPDH protein expression in wild-type and *Sharpin^cpdm^* MSFs plated in full medium on collagen I-coated 2 kPa PAA hydrogel or plastic directly after isolation from the mouse mammary gland. (**C**) Quantification of the relative integrin α11 expression levels detected by Western blot analysis of integrin α11 protein expression in wild-type and *Sharpin^cpdm^*MSFs cultured without serum and plated on collagen I., n_wt and Sharpincpdm_=8 samples from 4 independent experiments. GAPDH was detected for loading control. Mann–Whitney *U*-test, Mean ± SEM shown. (**D**) Validation of *Itga1* (upper panel) and *Itga11* (lower panel) silencing with siRNA by qPCR (expression normalized to GAPDH housekeeping gene), n=1 experiment.

**Table S1.**

| Parameter | meaning | Value | Origin |
| --- | --- | --- | --- |
| $N_{cl}$ | Number of FN molecules | 2000 | Adjusted |
| $K_{ont}$ | True binding rate | $0.1 \times 10^{-4}$ $\mu m^2/s$ wt<br>$0.8 \times 10^{-4}$ $\mu m^2/s$ cpdm | Adjusted, of the order of values reported for $\alpha IIB\beta 3$ (Litvinov et al., 2012) |
| $n_m$ | Number of myosin motors | From 2000 at 300 Pa to 18717 at 42129 Pa | Adjusted |
| $F_m$ | *Myosin motor stall force* | 2 pN | (Molloy et al., 1995) |
| $v_u$ | Unloaded myosin motor velocity | 110 nm/s | (Chan and Odde, 2008; Elosegui-Artola et al., 2014) |
| $d_{int}$ | Integrin density on the membrane | 1600 /$\mu m^2$ | Set, of the order of reported values (Elosegui-Artola et al., 2014) |
| $K_{off}$ | Integrin unbinding rate, scaling factor applied to force curve reported in Kong et al. | 0.05 wt<br>0.4 cpdm | Adjusted, catch bond dependency form (Elosegui-Artola et al., 2016; Kong et al., 2009) |
| $k_c$ | Clutch spring constant | 1 nN/nm | (Roca-Cusachs et al., 2012) |
| $r_a$ | Radius of adhesion | 1700 nm | Set |

